# Thermal acclimation of spreading depolarization in the CNS of *Drosophila melanogaster*

**DOI:** 10.1101/2024.05.31.596768

**Authors:** Mads Kuhlmann Andersen, R. Meldrum Robertson, Heath A. MacMillan

## Abstract

During exposure to extreme stress, the CNS of mammals and insects fails through a phenomenon known as spreading depolarization (SD). SD is characterized by an abrupt disruption of ion gradients across neural and glial membranes that spreads through the CNS, silencing neural activity. In humans, SD is associated with neuropathological conditions like migraine and stroke. In insects, it is coincident with critical thermal limits for activity and can be conveniently monitored by observing the transperineurial potential (TPP). We used the TPP to explore the temperature-dependence and plasticity of SD thresholds and SD-induced changes to the TPP in fruit flies (*Drosophila melanogaster*) acclimated to different temperatures. Specifically, we characterized the effects of thermal acclimation on the TPP characteristics of cold-induced SD, after which we induced SD *via* anoxia at different temperatures in both acclimation groups to examine the interactive effects of temperature and acclimation status. Lastly, we investigated these effects on the rate of SD propagation across the fruit fly CNS. Cold acclimation enhanced resistance to both cold- and anoxic SD and our TPP measurements revealed independent and interactive effects of temperature and acclimation on the TPP and SD propagation. This suggests thermodynamic processes and physiological mechanisms interact to modulate the threshold for activity through SD and its electrophysiological phenomenology. These findings are discussed in relation to conceptual models for SD and established mechanisms for variation in the thermal threshold for SD.

## Introduction

Given a sufficient dose abiotic stress, the central nervous system (CNS) of many animals, notably insects and mammals, will gradually lose function and shut down due to a physiological phenomenon known as spreading depolarization (1). Spreading depolarization (SD) is characterized by a rapid, local collapse of ion gradients across glial and neuronal cell membranes resulting in an almost complete cellular depolarization that spreads from its starting location into adjacent neural tissue where it silences electrical activity (2–5). Fortunately, SD is isolated to grey matter and neuropils and cannot spread across white matter or axon tracts (1). In mammalian and human physiology, SD mainly occurs as a consequence of pathological conditions (e.g. migraine and stroke, (6, 7)). In ectothermic animals, however, it is often found associated with abiotic stress tolerance. In insects, for example, SD limits organismal tolerance to heat, cold, and anoxia by inducing a state of neuromuscular paralysis (4, 8–11). SD similarly occurs at the limit for acute heat tolerance in fish (although the order of events is slightly different, see Andreassen et al. (12)).

Interestingly, these events are remarkably similar in terms of their electrical and physiological phenomenology in insects and vertebrates (13). Thus, while the CNS of insects is considered simpler than that of mammals, there is merit in using insect SD as a model for mammalian neuropathophysiology (1).

At critical limits of temperature tolerance, SD in the CNS renders insects comatose (10, 14). At low temperatures, this phenotype is often referred to as chill coma (15, 16), and the temperature where this occurs is often used as an estimate of the critical thermal minimum (i.e. CT_min_) (17–20). However, while SD and chill coma represent absolute limits to behaviour and fitness (i.e. an insect cannot walk, fly or reproduce while in SD-induced coma), the temperature at which these events occur is highly plastic and can vary substantially within and between insect species (21, 22). This is also true for the genus *Drosophila*, where these lower thermal limits can vary by more than 10°C among species held under common garden conditions (23, 24), and substantial variation can be elicited within a single species through thermal acclimation (25–27). Indeed, *Drosophila melanogaster* possesses an impressive capacity for thermal acclimation and differences of 8-12°C in CT_min_ have been observed in flies originating from the same population (25, 27, 28). Not surprisingly, the same trend has been observed for the temperature leading to cold-induced SD, which also varies between species and exhibits a similarly strong acclimation response (8, 17, 29). Thus, to gain a deeper understanding of the variation in the lower thermal limits in insects, we inevitably need to understand the processes that lead to variation in SD thresholds.

The physiological mechanism that triggers SD remains unknown and a topic of debate (see (1, 3, 30)), but it clearly leads to large, rapid changes to the distribution of ions across neuronal and glial membranes within the CNS (4, 5). This redistribution of ions is characterized by a surge in interstitial K^+^ concentration and the near-complete depolarization of neuronal membranes. It also depolarizes the adglial membrane of perineurial glia, and spreads from the affected area, invading previously unaffected tissue (1, 2). This adglial depolarization can be conveniently measured in insect preparations by using glass microelectrodes to penetrate the glial blood-brain barrier and monitor its transepithelial potential. This electrical potential is commonly referred to as the transperineurial potential (TPP) and is defined as the difference between the basolateral and adglial membrane potentials of the perineurial glia (V_b_ and V_a_, respectively, and TPP = V_b_ – V_a_; see (31)). Thus, when the adglial membrane depolarizes towards zero during the onset of a SD the TPP shifts in the negative direction with an amplitude that reflects V_a_ (see **Fig. 1**, and Robertson et al. (1)).

**Figure 1.**
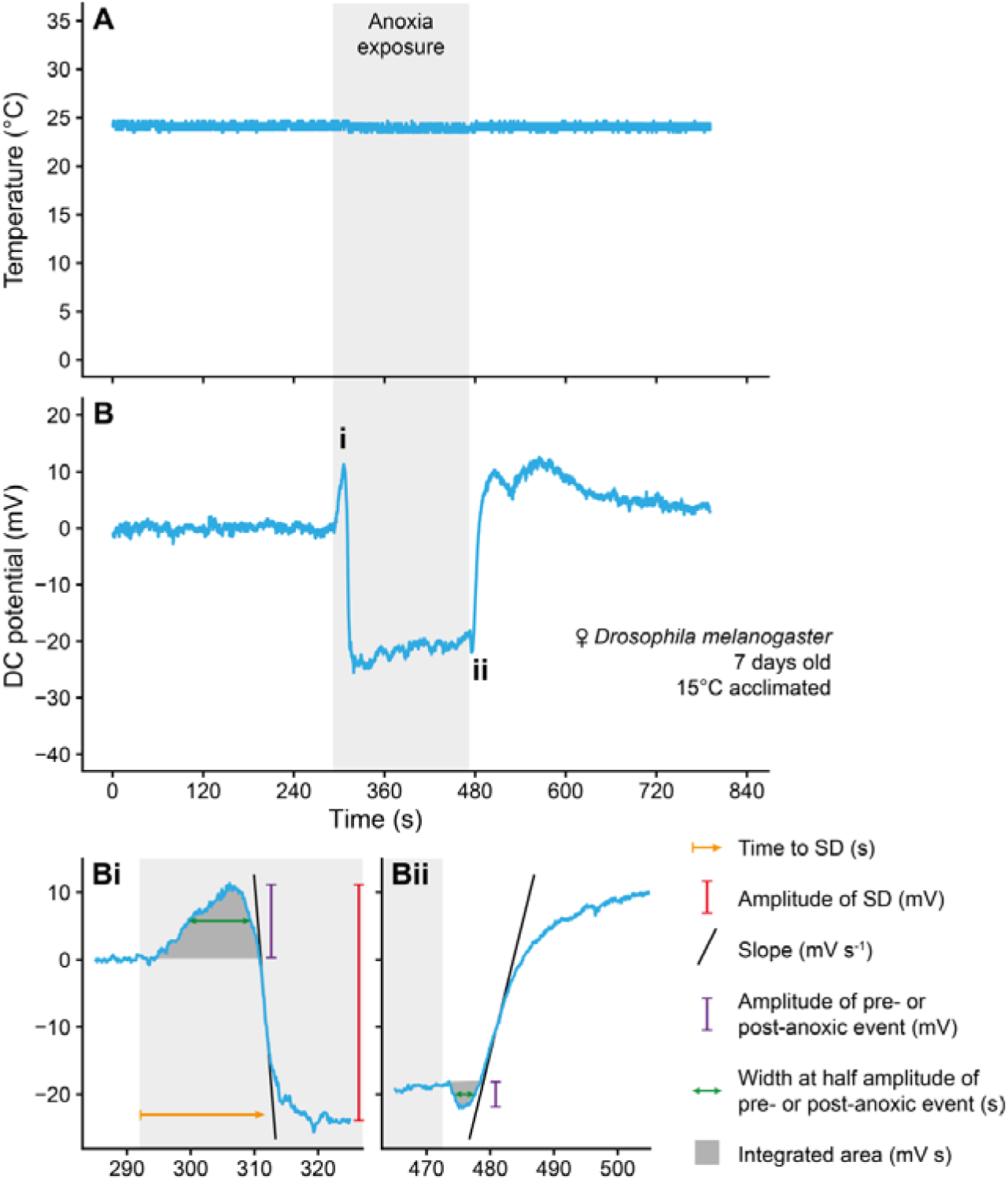
Example trace of the transperineurial potential (TPP) during an anoxia-induced spreading depolarization. In experiments where the effects of temperature and thermal acclimation on the spreading depolarization TPP waveform were investigated using anoxia we obtained matching measurements of (A) temperature and (B) transperineurial potential (which we refer to as a DC (direct current) potential here as it has been zeroed at the beginning of the trace). Below panel B are magnified views of (Bi) the pre-anoxic positivity and the negative shift in TPP caused by the spreading depolarization and (Bii) the post-anoxic negativity followed by the slow recovery of TPP after reoxygenation. Exposure to anoxia is marked in a light grey, shaded area. From these experiments we parameterized the waveform by quantifying 1) the time it took for the anoxia exposure to induce a spreading depolarization event (orange), 2) the amplitude of the shift in TPP caused by the event (red), 3) the slope of the TPP during the onset and recovery from the anoxic spreading depolarization (black), 4) the amplitude of the pre-anoxic positivity and post-anoxic negativity (purple), 5) the width at the half-amplitude of the pre-anoxic positivity and post-anoxic positivity (green), 6) the integrated area under the curve of the small pre- or post-anoxic shifts in TPP (dark grey). Note that the switch to anoxia often caused a minor change in temperature (see Fig. S1).

This size of the negative TPP shift is often referred to as the SD amplitude (or TPP amplitude) and the shift generally takes 5-10 s from beginning to completion, depending on the stressor responsible for the event (e.g. temperature, anoxia). From here, the TPP will stay in this state until the abiotic stressor is removed, after which TPP (and the insect) recovers, albeit at a slower rate than during SD onset (provided the stressor did not cause injury; (9, 13)). Recovery is often followed by a transient overshoot in TPP after which it returns to baseline. In the case of anoxia-induced SD there is also a small positive shift in TPP immediately before the SD and a small negative shift immediately before the recovery of the TPP. Combined, these parameters make up and characterize the TPP waveform of the SD and can be used to infer changes regarding the integrity of the glial blood-brain barrier, interstitial ion concentrations, and adaptive changes to electrogenic transporters (10, 32, 33).

The TPP waveform or an equivalent electrophysiological measurement (e.g. negative shift in the extracellular direct current potential), has been used to signal the onset of (and recovery from) SD in mammals, fish, and insects (9, 11, 13, 17, 34–36). Similarly, features of the TPP waveform have been used to infer changes to physiological function in comparative studies. For example, cold acclimation results in a more positive TPP at room temperature and reduces the TPP amplitude during cold-induced SD in *D. melanogaster*, suggesting adaptive changes to Na^+^/K^+^-ATPase localization and thermal sensitivity (29, 37). That being said, such inference can be potentially misleading as epithelial potentials are directly affected by temperature; it is a factor in empirical models of membrane potentials [e.g. the Goldman-Hodgkin-Katz equation (38, 39) and the charge difference model ((40), see Bayley et al. (41) for its application to insects)]. Thus, if SDs happen at different temperatures in a comparative model system, temperature itself may confound interpretations of differences in the TPP waveform. Similarly, the amplitude of the TPP waveform during temperature-induced SD is unlikely to reflect a baseline/control V_a_ under such conditions.

Recent studies have linked specific physiological functions to certain TPP waveform parameters. As mentioned, the TPP amplitude (**Fig. 1B**) is thought to be a measure of V_a_ immediately before the SD (1). The small positive shift before the abrupt TPP drop (**Fig. 1Bi**) has been hypothesized to represent the activity of electrogenic ion-motive pumps (42). Lastly, the small negative shift immediately before recovery (**Fig. 1Bii**) has been linked to reactivation of the V-type H^+^ ATPase to re-establish disrupted H^+^ gradients, while the slope of TPP recovery is reduced by inhibiting the Na^+^/K^+^-ATPase (43). Thus, quantifying the effects of temperature and thermal acclimation on the TPP waveform can provide interesting, albeit indirect, insight into underlying physiological or molecular mechanisms.

Much like aspects of the TPP waveform have been used to generate and test mechanistic hypotheses, so has the ability for the event to spread from acutely affected neural tissue to adjacent naïve tissue (2, 5, 13). The rate of propagation is generally described as ranging from ∼ 2-3 mm min^-1^ in insects (42, 44) and ∼ 2-9 mm min^-1^ in mammals (13, 45). However, endotherms (e.g. mammals and birds) and ectotherms (e.g. insects) often operate at vastly different body temperatures, making direct comparisons inadvisable. Indeed, the propagation velocity has been shown to be temperature-sensitive with an exponential increase up to temperature where propagation velocity was maximized followed by a sharp decline at supraoptimal temperatures (46). Thus, to accurately compare information collected on different model systems, temperature effects need to be accounted for (3).

To substantiate mechanisms of thermal acclimation it is critical to characterize and discount the direct effects of temperature on SD parameters. Such an approach allows us to tease apart thermodynamic effects from those caused by physiological and molecular mechanisms of thermal acclimation in the CNS. Here, we acclimated *Drosophila melanogaster* to relatively cold and warm temperatures throughout development and performed three experiments: First, we induced SD with stressful cold to characterize the effects of acclimation on the TPP waveform. Second, to investigate if the observed effects of acclimation were caused by the SD occurring at different temperatures or the acclimation treatment, we estimated the isolated effects of temperature by using anoxia to trigger SD events at different temperatures and characterizing differences in the waveform between acclimation groups. Lastly, we investigated the effects of temperature and thermal acclimation on the rate at which the SD propagates across neural tissue.

## Materials and Methods

### Animal maintenance and acclimation treatment

Our population of *Drosophila melanogaster* (Meigen, 1830) was established from isofemale lines of flies collected from London and Niagara on the Lake (both Ontario, Canada) in 2007 (47). The population was reared in 200 mL plastic bottles with ∼ 40 mL of food (see recipe in (29)) and kept at 20°C with a 12:12 light cycle. Experimental flies were produced by allowing flies from the 20°C population to oviposit on fresh medium in a new bottle for 4-6 h to ensure a density of 150-200 flies per bottle. Egg-containing bottles were placed in one of two temperature-controlled incubators (MIR-154-PA, Panasonic) for developmental acclimation to either 15 or 25°C (12:12 light cycle).

Bottles were checked daily and newly emerged flies were transferred to 40 mL glass vials with ∼ 7 mL of the same food and left to mature for 6-8 days at their acclimation temperatures. To avoid sex-specific differences, only females were used in experiments.

### Experimental preparation and electrophysiology

Glass microelectrodes were pulled from borosilicate glass (1B100F-4, WPI, Sarasota, FL, USA) on a Flaming-Brown type micropipette puller (P-1000, Sutter Instruments, Novato, CA, USA) to a tip resistance of 5-7 MΩ when backfilled with a 500 mM KCl solution. This electrode was placed in a half-cell electrode holder with a Ag/AgCl wire, attached to a micromanipulator (M3301-M3-L, WPI), and connected to a Duo 773 intracellular/extracellular electrometer (WPI). Raw voltages were digitized by a PowerLab 4SP A/D converter (ADInstruments, Colorado Springs, CO, USA) and fed to a computer running LabChart 4.0 software (ADInstruments).

Electrophysiological measurements of SD were made in individual, restrained flies. To avoid confounding effects of anesthesia (e.g. (48)), an aspirator with a 100 mL pipette tip attached to the end was used to manipulate individual flies by their heads after which they were secured in wax on a glass cover slide and prepared as described by Spong, Rodríguez and Robertson (42). Here, a pair of micro-scissors was used to cut two holes in the fly: one small hole immediately above the ocelli, and another in the abdomen. The fly preparation was then moved to a custom-built, thermoelectrically-cooled stage under a microscope. Here, the glass microelectrode was inserted into the hemolymph surrounding the brain while a ground Ag/AgCl wire was inserted into the abdomen. Once the fly was prepared for electrophysiology, a type K thermocouple was placed next to the head at the same height above the cooling stage to monitor ganglion temperature. This was connected to a channel of the A/D converter with a thermocouple-to-analog converter (TAC80B-K; Omega, Stamford, CT, USA). The glass electrode was then slowly inserted into the extracellular space of the brain using the micromanipulator. During this insertion, the measured potential should show a small, sudden increase indicative of the blood-brain barrier having been penetrated, and the magnitude of this shift represents the transperineurial potential (TPP). From here, three different experimental procedures were followed as outlined below.

### Experiment 1 – Cold-induced spreading depolarization

First, SD was induced with stressful cold. For this experiment, the TPP at room temperature (∼ 23°C) was recorded after which the temperature was decreased at a rate of ∼ 1°C min^-1^ (according to the thermocouple). The temperature was lowered until a rapid drop in TPP of 25-50 mV and ∼ 5-10 s in duration was observed, which is indicative of a SD having occurred (42). The temperature at which the event occurred was measured as the temperature at half the amplitude of the negative shift in TPP. After cooling for another 1 min, the preparation was brought back to room temperature to observe the recovery of TPP. Apart from the onset temperature a number of parameters relating to the SD waveform were characterized: 1) amplitude of TPP drop (in mV), 2) slope of the steepest part of the drop in TPP (onset slope; in mV s^-1^), 3) slope of the steepest part of the recovery of TPP during rewarming (recovery slope, in mV s^-1^), and 4) the temperature at which the recovery slope was estimated (in °C) (49). This experiment was performed on eight flies from each acclimation group (total N = 16).

### Experiment 2 – Effects of temperature on the anoxic spreading depolarization waveform

The second experiment was aimed at describing the isolated effect(s) of temperature on the previously noted changes in the TPP waveform during SD. To do so we induced SD with anoxia at different temperatures and characterized the TPP waveform. To avoid confounding effects of air flow, the experiment was set up such that the fly preparation was continuously exposed to a flow of atmospheric air (generated by a small aquarium pump) which could be changed to a flow of pure, compressed N_2_ gas *via* a 3-way valve (similar to Rodríguez and Robertson (50)). This valve was placed before a 1/8” (3.175 mm) Nalgene rubber tube (ThermoFisher Scientific, Waltham, MA, USA), through which the air flow was adjusted to ∼ 0.8 L min^-1^ through a series of clamps. This tube directed the gas flow to the preparation from a distance of ∼ 1.5 cm. Both air and N_2_ gas were sent through humidifiers before reaching the fly to avoid any risk of desiccation.

The flies were prepared as described above, after which the preparation temperature was set between ∼ 11 and ∼ 35°C. As temperature changed to reach the setpoint, the TPP was continuously monitored to avoid accidental temperature-induced SDs. After reaching the target temperature, the preparation was held at that temperature for 5 min to ensure that the SD was not induced. We found that temperatures ∼ 5°C above the temperature of cold-induced SD were safe for both warm- and cold-acclimated flies. Once the preparation was deemed stable, the gas was changed from air to N_2_ to trigger a SD. Exposure to N_2_ was maintained for 3 min after which air was reintroduced to monitor the recovery. From these experiments, the following anoxic SD waveform parameters were quantified: 1) time to SD after beginning of anoxia exposure (in s), 2) amplitude of drop in TPP (in mV), 3) onset slope (in mV s^-1^), and 4) recovery slope (in mV s^-1^). Furthermore, the TPP waveform elicited by an anoxic SD displays a characteristic positive shift before the big drop in TPP and a negative shift in potential before the TPP recovery phase (42). These two events, referred to as the pre-anoxic positivity and post-anoxic negativity, respectively, were analyzed by measuring their 1) amplitude (in mV), 2) width (at half-amplitude, in s), and integrated area (area above/below the TPP immediately before the post/pre-anoxic event, in mV s). The shift from air to N_2_ sometimes resulted in a small change in temperature (mean = -0.27°C, standard deviation = 1.04°C, range = -3.38 to 1.30°C) and was always in the direction of room temperature (see **Fig. S1**). Thus, the experimental temperature was quantified as the temperature at the time of the SD (as described for cold-induced SD). An example trace can be seen in **Fig. 1**. Experiments were performed on 30 flies from each acclimation group (total N = 60).

### Experiment 3 – Effects of temperature on the velocity of the spreading depolarization

To estimate the velocity at which the SD propagates across the brain, we used a modified version of the setup described above which incorporated a second glass microelectrode (42). Specifically, the second electrode, connected to a Neuroprobe amplifier (model 1600, A-M Systems, Sequim, WA, USA), was inserted into the brain at a distance away from the first electrode. After insertion of both electrodes, a Nanoject II injector (Drummond Scientific Company, Broomal, PA, USA) was used to inject 2-3 nL aliquots of 150 mM KCl solution into the head capsule. The KCl solution was injected at the edge of the opening in the head to artificially trigger a SD at one side to have it propagate across the brain and be observed by both electrodes. After the experiment, the electrodes were retracted, the fly removed, and the electrodes moved back to their original position by use of the micro-calipers so that the distance between the electrodes could be measured with a reticle in the microscope. The spreading velocity (in mm min^-1^) was then calculated based on the distance between the electrodes (in mm) and the delay in the negative shift in TPP between the two electrodes (in min). An example trace can be seen in **Fig. 2**. This experiment was performed at three temperatures (15, 23, and 30°C) in seven cold- and six warm-acclimated flies at each temperature (total N = 39).

**Figure 2.**
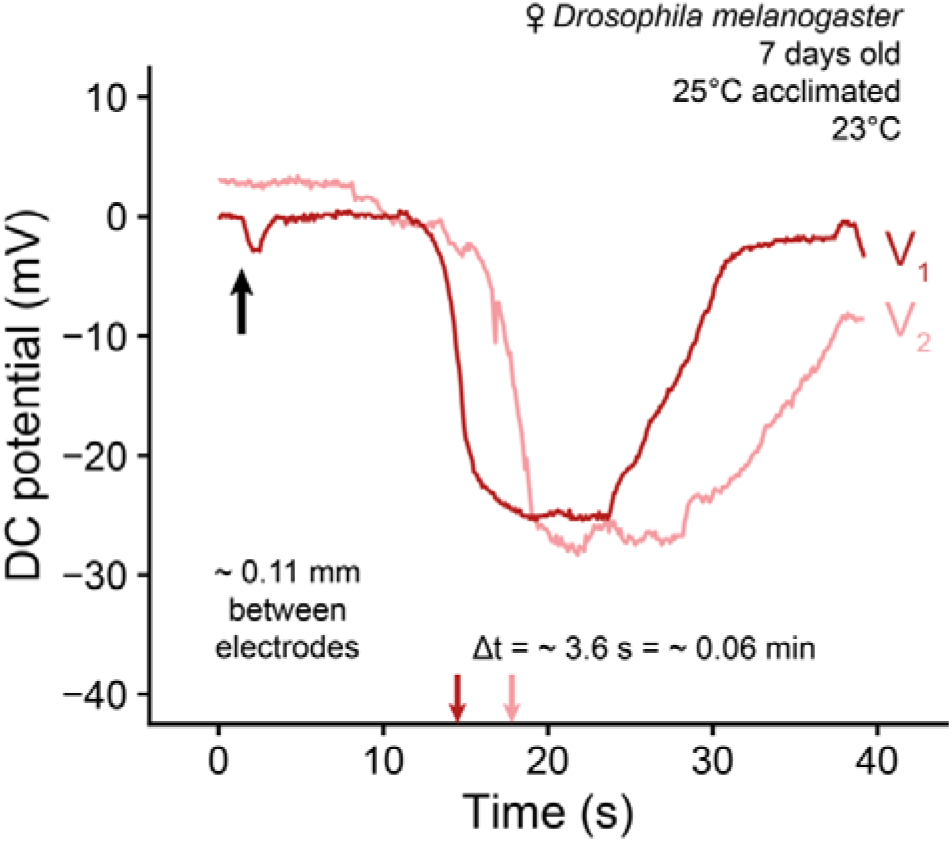
Example trace of the transperineurial potential measured at two sites with a spreading depolarization being triggered by KCl injection. Depicted is an example trace where KCl was injected into the head of a fly (black arrow and small artifact) which triggered a spreading depolarization after a short delay. This spreading depolarization propagated from one microelectrode (V_1_, dark red) to another placed ∼ 0.11 mm away (V_2_, light red) after a ∼ 3.6 s delay (dark- and light red arrows), resulting in a propagation velocity of ∼ 1.8 mm min^-1^. The potential traces have been zeroed and shifted vertically for clarity, meaning that they no longer represent the TPP and we therefore refer to them as DC potentials.

### Statistical analysis

All data analysis was performed in R software (version 4.2.2 (51)). The three datasets were analyzed as follows:

*Experiment 1*: Parameters for cold-induced SD were first tested for normality using Shapiro-Wilk’s tests. One dataset (SD recovery slope) violated normality and acclimation groups were therefore compared with a Mann-Whitney-Wilcoxon test. For remaining parameters, group variances were compared with F tests. Datasets with similar group variances were compared with Student’s t tests while Welch t tests were used for those with dissimilar variances.

*Experiment 2*: Effects of temperature on the parameters associated with the TPP waveform during anoxic SDs were analyzed with either a linear model or non-linear regression. For linear models, the interaction term (temperature-acclimation interaction) was excluded from analyses is this was non-significant. Non-linear regression was performed using the nls() function and on only one parameter; the recovery slope. Here, a Gaussian model was fit to the dataset and the effect of acclimation was investigated by including an interaction term in the temperature variable (see (37)). Lastly, because the pre-anoxic positivity and post-anoxic negativity have been shown to relate directly to ion transport function we did an Arrhenius transformation followed by a linear regression. This transformation fit the data well as it improved the AIC score substantially (125.0 to 96.6) and reduced residual variance (by ∼ 44% from 0.42 to 0.24).

*Experiment 3*: The relationship between temperature and propagation of the KCl-induced SD was exponential, and the effects of temperature and acclimation were analyzed using a linear model on log-transformed propagation velocities. Q_10_ values were calculated using the Q10() function from the *respirometry* package (52).

For all analyses, the level for statistical significance was set to 0.05. All values presented are mean ± standard error of the mean unless otherwise stated.

## Results

### Cold-acclimation lowers the temperature for cold-induced spreading depolarization and alters the associated TPP waveform

Acclimation temperature greatly affected the temperature leading to SD (t_14_ = -7.2, P < 0.001) such that cold-acclimated flies had a lower threshold for cold-induced SD (5.5 ± 0.3°C) compared to their warm-acclimated conspecifics (9.9 ± 0.5°C; **Fig. 3A**).

**Figure 3.**
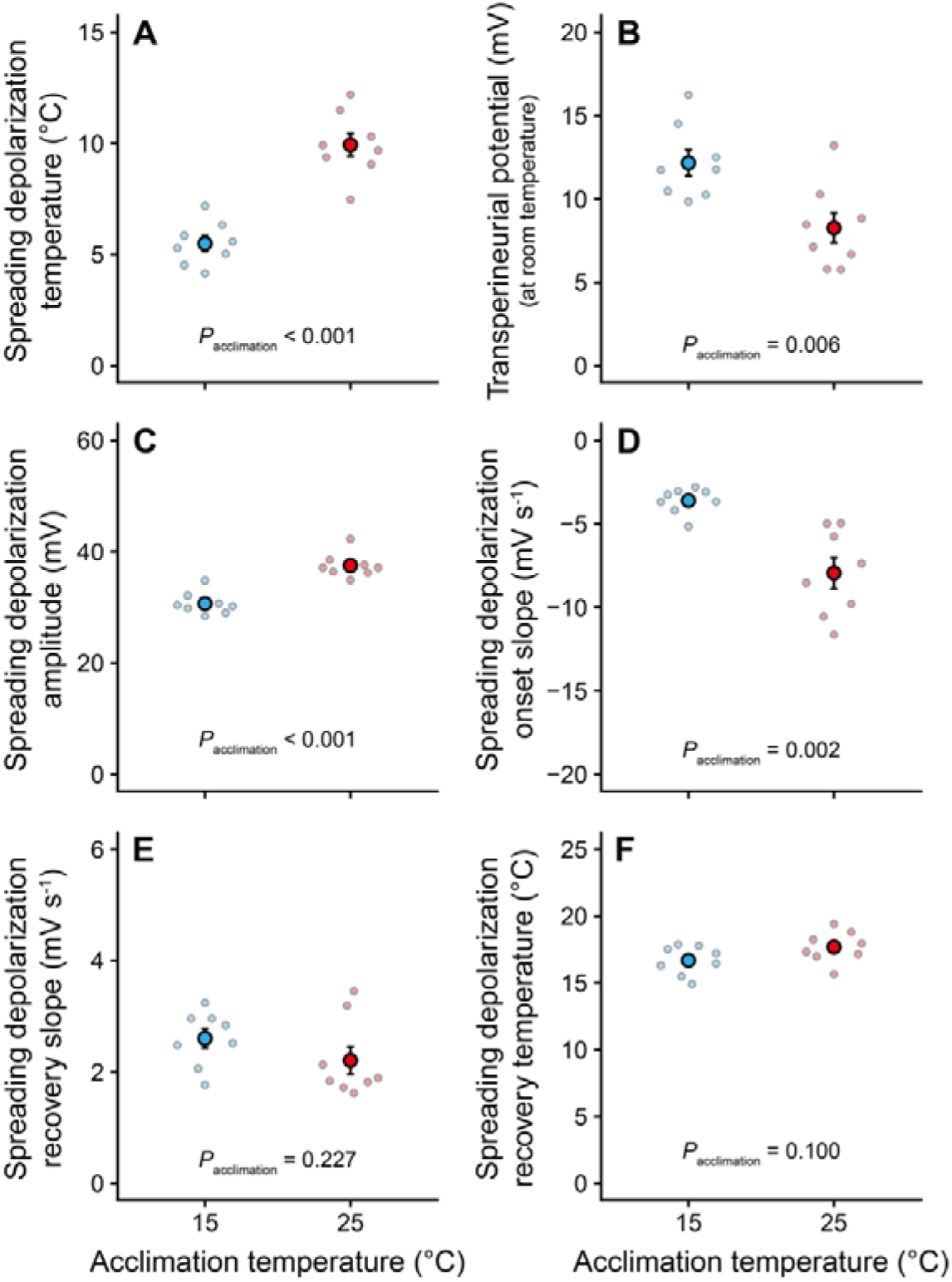
Effects of thermal acclimation on the temperature leading to cold-induced spreading depolarization, the transperineurial potential, and the spreading depolarization waveform. By monitoring the transperineurial potential during a temperature ramp from room temperature in both cold- and warm-acclimated flies (blue and red, respectively), we obtained not only the temperature leading to cold-induced spreading depolarization (A), but also the baseline transperineurial potential (B), the amplitude of the spreading depolarization (C), the slope of the TPP during the spreading depolarization onset (D), the recovery of TPP during rewarming (E), and the temperature at which the recovery slope was estimated (F). Small, scattered points depict individual measurements while the larger points denote the mean. Error bars not shown are obscured by the symbols.

Cold acclimation also led to a number of changes to the TPP and the SD-induced TPP waveform: TPP at room temperature was more positive after cold acclimation (12.2 ± 0.8 mV and 8.3 ± 0.9 mV for cold- and warm-acclimated flies, respectively, **Fig. 3B**; t_14_ = 3.3, P = 0.006), however, despite having a more positive starting TPP, cold-acclimated flies had a lower SD amplitude (30.7 ± 0.7 mV) compared to warm-acclimated flies (37.5 ± 0.8 mV) (**Fig. 3C**, t_14_ = -6.5, P < 0.001). Similarly, the cold-acclimated flies had a less steep slope in TPP during the SD than the warm-acclimated flies (-3.6 ± 0.3 mV s^-1^ and -7.9 ± 0.9 mV s^-1^, respectively, **Fig. 3D**; Welch t_8.2_ = 4.5, P = 0.002). Lastly, the slope of TPP recovery during rewarming did not differ between cold- and warm-acclimated flies (2.6 ± 0.2 mV s^-1^ and 2.2 ± 0.2 mV s^-1^, respectively, **Fig. 3E**; Mann-Whitney-Wilcoxon U = 44, N = 16, P = 0.227), and occurred at similar temperatures (16.7 ± 0.4°C and 17.7 ± 0.4°C, respectively, **Fig. 3F**; t_14_ = -1.8, P = 0.100).

### Temperature and thermal acclimation alter the anoxic spreading depolarization waveform

To investigate whether the differences above were driven by acclimation, effects of the different temperatures at which the SD occurred, or a combination thereof, we used anoxia to trigger SDs at different temperatures in both cold- and warm-acclimated flies.

The experimental approach allowed us to investigate the effects of temperature and thermal acclimation on the timing of the anoxic SD (example shown in **Fig. 1**), which can be interpreted as a measure of anoxia tolerance (**Fig. 4**). Anoxia tolerance was found to be temperature-sensitive (F_1,56_ = 198.8, P < 0.001), such that lower temperatures increased the time until the anoxia elicited a SD from ∼ 10 s at 35°C to more than 20 s at 15°C and below (i.e. improved anoxia tolerance). At the same time, it took an average of 9.4 ± 2.9 s longer for the anoxia to cause SD in cold-acclimated flies compared to warm-acclimated flies (F_1,56_ = 19.2, P < 0.001). These effects were synergistic such that the effect of temperature was greater in cold-acclimated flies, which generally manifested as longer exposures to anoxia before SD at lower temperatures (e.g. ∼ 30 s and ∼ 22 s at 15°C for cold- and warm-acclimated flies, respectively; interaction: F_1,56_ = 4.4, P = 0.040) – a trend which continued for cold-acclimated flies below the lowest temperature where we were able to obtain reliable measurements for warm-acclimated flies (i.e. ∼ 32 s at 11°C).

**Figure 4.**
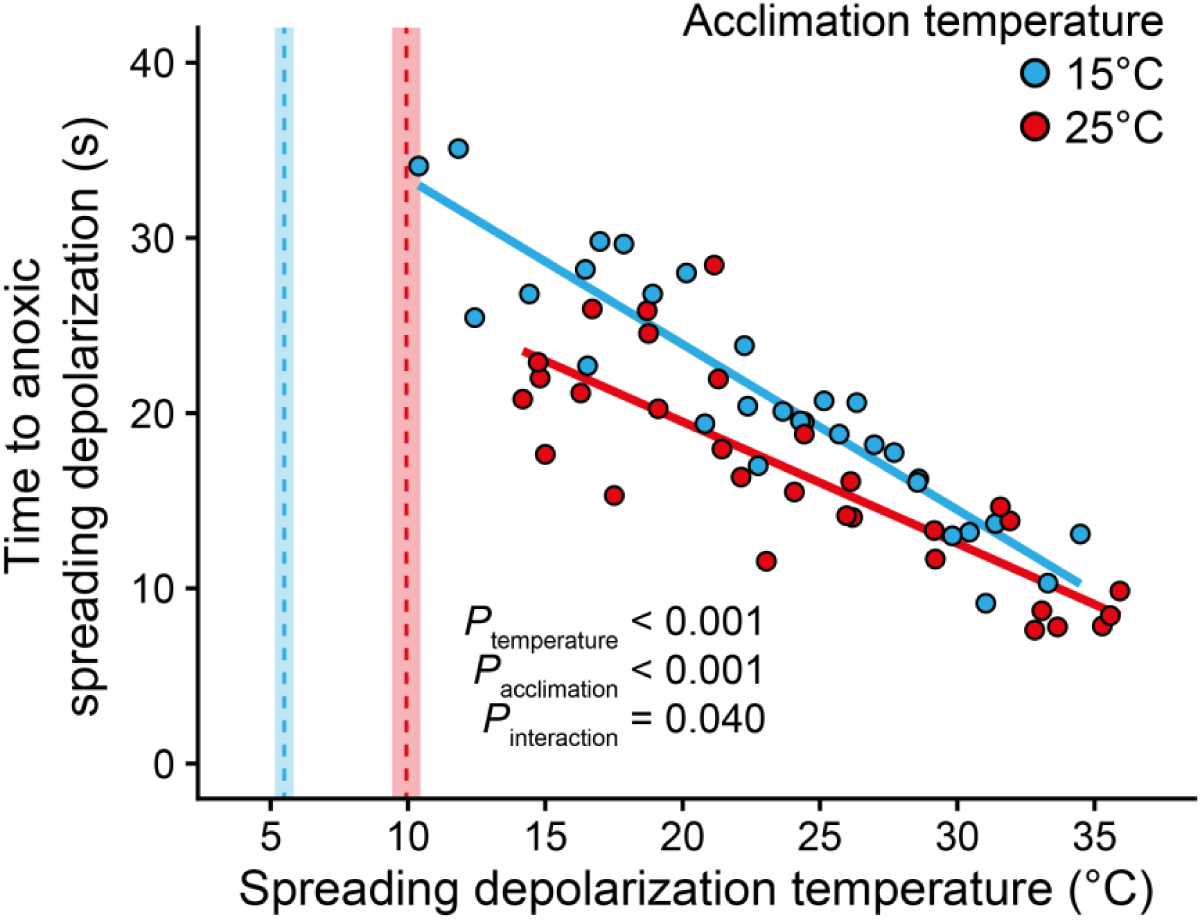
Thermal acclimation and temperature interact to modulate sensitivity of the brain to anoxia. The time until a spreading depolarization occurred, quantified as the time from the beginning of anoxia exposure to the midpoint in the drop in TPP, was used as an estimate of sensitivity to anoxia. Cold- and warm-acclimated flies are depicted in blue and red, respectively, with their linear regressions. Vertical dashed lines and shaded areas denote the mean temperature and standard error, respectively, where spreading depolarization is caused by the low temperature itself (see **Fig. 3A**).

After investigating the timing of the anoxic SD as a measure of anoxia tolerance we parameterized the associated TPP waveform (as shown in **Fig. 1**) by quantifying the large shifts in TPP that denote the onset and recovery from the SD (**Fig. 5**). Furthermore, we quantified the smaller pre- and post-anoxic events to gain insight into putative physiological changes induced by temperature and acclimation (**Fig. 6**).

**Figure 5.**
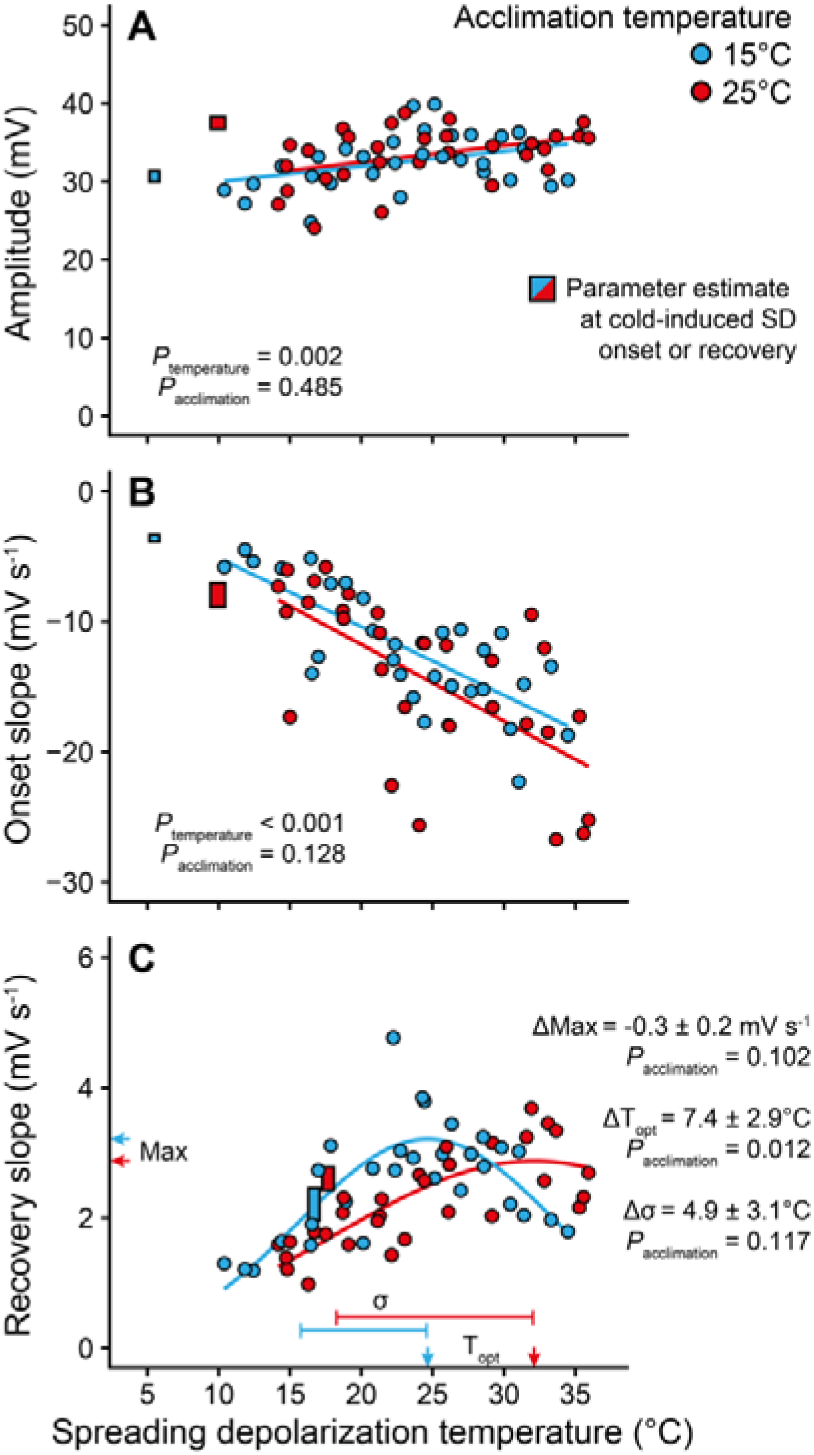
Major shifts in the transperineurial potential that denote the onset of spreading depolarization are temperature-sensitive and recovery is temperature- and acclimation-specific. During the onset of and recovery from spreading depolarization there are large shifts in TPP. We have quantified these changes in both cold- and warm-acclimated flies (blue and red, respectively) at temperatures ranging from 11 to 35°C and report these shifts as the (A) amplitude of the shift, (B) slope of descend during onset, and (C) slope of recovery after reoxygenation. Included in each panel are the linear or non-linear models that best predict the effects of temperature (blue and red lines for cold- and warm-acclimated flies, respectively). The non-linear model fit used to model the recovery slope is a Gaussian fit (see Andersen, Robertson and MacMillan (37)), which shows that the thermal performance curve for recovery has been left-shifted in the cold-acclimated group compared to the warm-acclimated group (i.e. ΔT_opt_). Each panel also contains the estimates obtained during the onset (i.e. amplitude and onset slope) and recovery (i.e. recovery slope) from cold-induced spreading depolarization and temperature of recovery therefrom obtained in *Experiment 1* (squares, blue and red for cold- and warm-acclimated flies). Note that these estimates for cold-induced spreading depolarization fit the trend of temperature-sensitivity in anoxic spreading depolarization well, except in the case of amplitude and recovery slope for the warm-acclimated group (**A**), indicating that anoxia and stressful cold might lead to spreading depolarization via slightly different mechanisms.

**Figure 6.**
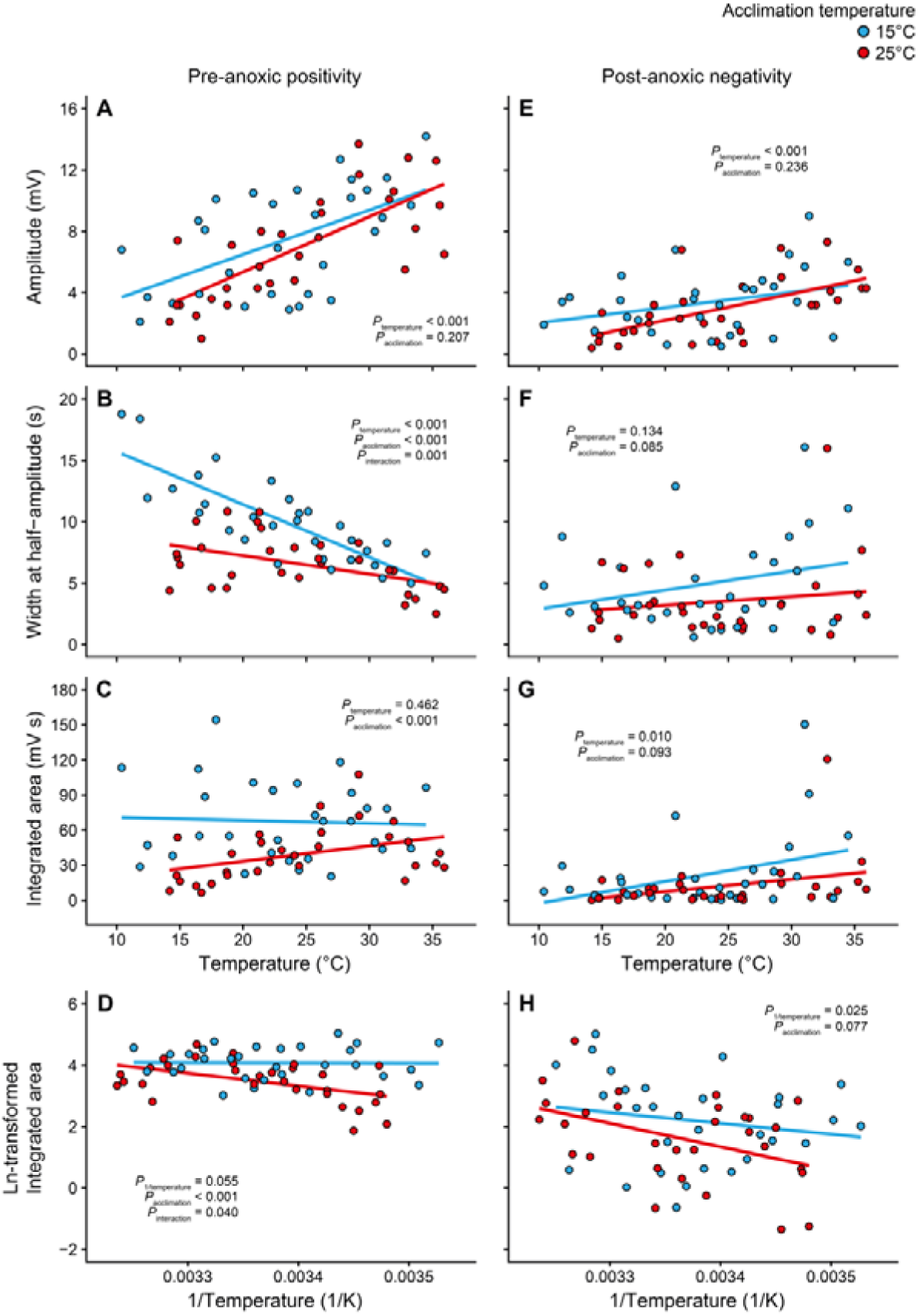
The pre-anoxic positivity is affected by temperature, acclimation, and their interaction while the post-anoxic negativity is temperature-sensitive. Before the major negative shift in TPP that denotes the onset of anoxia-induced spreading depolarization there is a smaller pre-anoxic positivity (left column), and immediately before the slightly slower positive shift in TPP that signifies recovery from spreading depolarization there is a small post-anoxic negativity immediately as oxygen is reintroduced (right column). These events were parameterized in both cold- and warm-acclimated flies (blue and red, respectively) by quantifying their amplitude (A and E), their duration at half-amplitude (B and F), and the integrated area under/over the curve (C and G). Lastly, we decided to Arrhenius-transform the integrated area as we speculate that these relate to changes in ion transport capacity (D and H). Linear models are depicted in each panel for each acclimation group. Note that these TPP parameters were not quantified for the cold-induced spreading depolarization as they were so small that they were often indistinguishable from noise in the electrophysiological trace.

The amplitude of the shift in the TPP during SD (**Fig. 5A**) increased with temperature (F_1,57_ = 11.0, P = 0.002) at a rate of ∼ 0.2 mV per 1°C from ∼ 30 mV at 11°C to ∼ 35 mV at 35°C, and this effect was independent of acclimation temperature (F_1,57_ = 0.5, P = 0.485). Similarly, we found that the onset slope in TPP during SD onset (**Fig. 5B**) was temperature-dependent (F_1,57_ = 55.2, P < 0.001) and increased in steepness by ∼ -0.6 mV s^-1^ per 1°C increase in temperature from ∼ -6.0 mV s^-1^ at 11°C to ∼ -20 mV s^-1^ at 35°C. Acclimation temperature did not affect the onset slope (F_1,57_ = 2.4, P = 0.128). The relationship between the slope of TPP recovery after the reintroduction of oxygen and temperature was clearly non-linear (**Fig. 5C**) so we decided to analyze this part of the dataset using non-linear regression to a Gaussian model following Andersen, Robertson and MacMillan (37). Here we found that the maximum recovery slope (ΔMax) and thermal sensitivity (Δσ) were unaffected by acclimation (t = -1.7, P = 0.102 and t = 1.6, P = 0.117, respectively, with N = 60 and df = 54). Conversely, the temperature where the recovery slope was optimized (ΔT_opt_) was reduced by 7.4 ± 2.9°C in the cold-acclimated flies compared to the warm-acclimated group (t = 2.6, P = 0.012 with N = 60 and df = 54).

A pre-anoxic positivity event was observed at all temperatures in both acclimation groups (**Fig. 6A-D**). The amplitude of the event increased with temperature (F_1,57_ = 40.3, P < 0.001) from ∼ 4 mV at 11°C to ∼ 10 mV at 35°C and was independent of acclimation (F_1,57_ = 1.6, P = 0.207) (**Fig. 6A**). The duration of the pre-anoxic positivity, estimated as the width at half-amplitude, was also temperature-sensitive and decreased with temperature (F_1,56_ = 59.6, P < 0.001). Interestingly, we found that cold-acclimation increased the duration of the pre-anoxic positivity (F_1,56_ = 35.7, P < 0.001) and that acclimation affected the temperature-sensitivity (interaction: F_1,56_ = 12.8, P = 0.001) (**Fig. 6B**). Specifically, the duration of the pre-anoxic positivity for both cold- and warm-acclimated flies was ∼ 5 s at 25°C while the event lasted longer for cold-acclimated flies at low temperatures (∼ 15 s at 11°C) compared to warm-acclimated flies (∼ 8 s at 11°C). When quantifying the overall, integrated area under the curve of the pre-anoxic positivity, we were surprised to find that this was unaffected by temperature (F_1,57_ = 0.5, P = 0.463) but elevated in the cold-acclimated group (F_1,57_ = 15.1, P < 0.001) (**Fig. 6C**). Under the expectation that the pre-anoxic positivity reflects an aspect of ion transport capacity (e.g. activation of the Na^+^/K^+^-ATPase as per Spong, Rodríguez and Robertson (42)) we decided to Arrhenius-transform the integrated area under the curve and repeat the analysis (**Fig. 6D**). Doing so, the pre-anoxic positivity remained unaffected by temperature (despite approaching statistical significance; F_1,56_ = 3.8, P = 0.055) while cold-acclimated flies still had a higher integrated area (F_1,56_ = 18.5, P < 0.001). Lastly, we found a significant interaction between temperature and acclimation (F_1,56_ = 4.4, P = 0.040) such that the event was the same size in both groups at high temperatures (left side in Arrhenius plot) and decreased in warm-acclimated flies at the lower temperatures (right side in Arrhenius plot) while it remained unchanged in the cold-acclimated group.

Like the pre-anoxic positivity, the post-anoxic negativity was observed at all temperatures in both acclimation groups, however, it was generally smaller in amplitude and shorter in duration (**Fig. 6E-H**). Note that while the negative shift in TPP during the post-anoxic negativity constitutes negative amplitudes and integrated areas, we will be referring to them in absolute terms for the sake of clarity. The amplitude of the post-anoxic negativity was affected by temperature (F_1,57_ = 15.0, P < 0.001), unaffected by acclimation (F_1,57_ = 1.4, P = 0.236), and ranged from ∼ 1-2 mV at the lower temperatures to ∼ 5 mV at 35°C (**Fig. 6E**). The duration of the event ranged from 0.5 to 16 s and was affected by neither temperature (F_1,57_ = 2.3, P = 0.134) nor acclimation (F_1,57_ = 3.1, P = 0.085) (**Fig. 6F**). This results in an integrated area that much like the amplitude is temperature-sensitive (F_1,57_ = 7.1, P = 0.010) and unaffected by acclimation (F_1,57_ = 2.9, P = 0.093) (**Fig. 6G**). Similarly to the pre-anoxic positivity, we Arrhenius-transformed this dataset (**Fig. 6H**), which revealed a positive effect of temperature on integrated area of the post-anoxic negativity (F_1,57_ = 5.3, P = 0.025) that was independent of acclimation temperature (F_1,57_ = 3.3, P = 0.077).

### Effects of temperature and thermal acclimation on the spreading depolarization propagation velocity

To further characterize the effects of temperature and thermal acclimation on SD parameters we quantified the velocity at which the SD propagated across the *D. melanogaster* brain (**Fig. 7**). We did so by triggering a SD with KCl and using two electrodes at different positions to capture the event as it spread (example trace in **Fig. 2**). We found a strong exponential effect of temperature on propagation velocity (F_1,35_ = 133.0, P < 0.001) and no overall effect of acclimation (F_1,35_ = 0.1, P = 0.788). Interestingly, acclimation temperature affected the thermal sensitivity of the propagation velocity (interaction: F_1,35_ = 4.5, P = 0.040) such that the SD propagated faster in cold-acclimated flies when measured at 15°C (1.2 ± 0.2 mm min^-1^ and 0.9 ± 0.2 mm min^-1^ for cold- and warm-acclimated flies, respectively) but slower when measured at 31°C (cold: 3.8 ± 0.1 mm min^-1^, warm: 5.0 ± 0.6 mm min^-1^). At room temperature the propagation velocities were similar (cold: 2.1 ± 0.2 mm min^-1^, warm: 2.5 ± 0.3 mm min^-1^). These differences in thermal sensitivity were also evident from the lower Q_10_ value for propagation velocity in the cold-acclimated flies (2.13) compared to warm-acclimated flies (2.98). Arrhenius transforming the dataset, we found similar effects of temperature (reciprocal of absolute temperature (1/K); F_1,35_ = 132.1, P < 0.001) and an interaction between acclimation and temperature (again 1/K) (F_1,35_ = 4.5, P = 0.042) with no effect of acclimation itself (F_1,35_ = 0.1, P = 0.773). Based on this transformation, the Arrhenius activation energy (E_a_) was calculated as being 55.9 kJ mol^-1^ (0.58 eV) and 80.8 kJ mol^-1^ (0.84 eV) for cold- and warm-acclimated flies, respectively.

**Figure 7.**
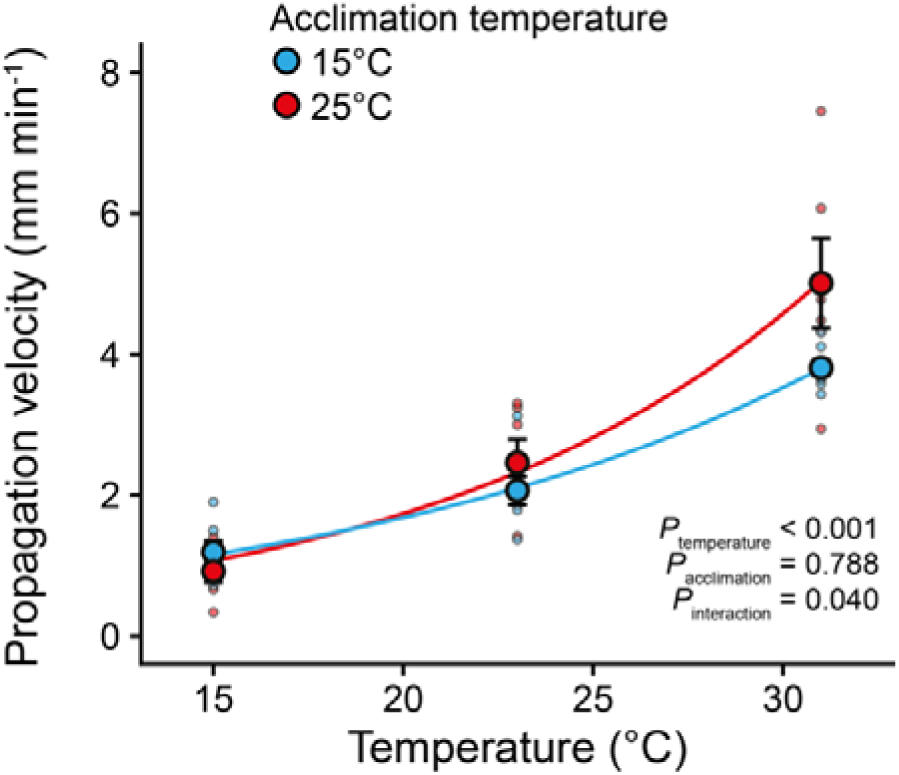
The relationship between the propagation velocity of KCl-induced spreading depolarization and temperature is exponential and it is affected by thermal acclimation. Propagation velocity was measured by triggering a spreading depolarization with KCl and monitoring the TPP at two different sites in the brain (example trace in **Fig. 2**). This experiment was repeated in both cold- and warm-acclimated flies (blue and red, respectively) at 15, 23, and 31°C to investigate the thermal sensitivity. Small translucent points depict individual measurements while larger opaque points denote the mean. Error bars not shown are obscured by the symbols.

## Discussion

When neural tissue is sufficiently challenged by an abiotic stressor it shuts down by a process known as a SD, which propagates across and through grey matter or neuropils as a wave of depolarization that arrests neuronal activity (1, 5). In mammals, this is generally associated with neuropathological conditions (3, 6, 53) and in insects it is most often observed at the critical thermal limits (8, 9, 17). Insects display impressive plasticity in the critical thermal limits (22), and these adaptive changes have been shown to relate to a variety of molecular, cellular, and physiological functions, many of which are shared with mammalian nervous system physiology (1, 10, 13). Thus, there is a growing interest in using insect SD as a translational model system to study the mechanisms underlying its initiation and variation of its threshold to improve human and mammalian neural health.

In insects, SD is often monitored by measuring the transperineurial potential (TPP), which has a characteristic waveform during the event (e.g. **Fig. 1**; see (1) and (42)). However, the properties of this waveform are not constant and are affected by a wide range of factors including thermal acclimation (see Andersen, Robertson and MacMillan (37) and the review by Robertson, MacMillan and Andersen (10)), which makes it challenging to directly compare waveforms from different acclimation groups. Furthermore, because the event itself occurs at different temperatures, the effect of acclimation cannot be separated from any direct effects of temperature. In the present study we aimed to disentangle the direct thermodynamic effects of temperature on the TPP waveform from those caused by acclimation by comparing waveforms at low temperature to similar waveforms elicited at different temperatures in cold- and warm-acclimated flies. We found that thermal acclimation alters the general stress tolerance of the CNS, and that acclimation status and temperature both contribute to properties of the TPP waveform in independent and interactive ways.

### Cold acclimation improves tolerance to cold and anoxia

Our acclimation treatment led a 0.44°C mean difference in the temperature leading to cold-induced SD per 1°C difference in acclimation temperature (**Fig. 3A**). This is similar to differences observed in previous studies on not only SD (∼0.3-0.4°C per 1°C difference) (17, 29, 37), but also the resulting organismal loss of function (∼0.4°C per 1°C difference) (25, 26, 28).

Anoxia tolerance, here quantified at the time to SD onset after the switch to a pure N_2_ atmosphere, was increased by both low temperature, cold acclimation, and their interaction. This is similar to the previously reported increased time to anoxia-induced SD in fruit flies exposed to mild cold (50), and improved anoxia tolerance of fruit flies with induced or adaptive improvements to cold tolerance (17). The protective effect of low temperatures on organismal anoxia tolerance has been well-studied, and is likely conferred through the associated cold-induced reduction in metabolic demands (54–57). Metabolic adjustments during cold acclimation are abundant (58, 59), however, overall metabolic rate has not been found to change through thermal acclimation or adaptation in fruit flies (60–62). Nonetheless, there is evidence to suggest that improved cold tolerance of the CNS relates to a reduced metabolic demand, reminiscent of channel- or spike arrest mechanisms (63–66). Specifically, cold-acclimated *D. melanogaster* have a generally lowered Na^+^/K^+^-ATPase activity in the brain and further exhibit a lower pump activity at the temperature leading to cold-induced SD (37). Similarly, improved cold tolerance in migratory locusts (*Locusta migratoria*) is associated with K^+^ sensitivity of the BBB (33) suggesting a reduced K^+^ conductance of the basolateral membrane (67).

### Concurrent effects of temperature and thermal acclimation alter the timing and waveform of spreading depolarization

After establishing a basis for studying the effects of thermal acclimation and temperature, we moved on to characterizing and comparing TPP waveforms. Here too, we replicated some effects described in previous studies (see **Fig. 3**); an increase in room temperature TPP after cold acclimation by ∼ 4 mV (29), and a reduced TPP amplitude (∼ 7 mV; ∼ 1.5 mV per 1°C change in temperature of SD) and onset slope (∼ 4-5 mV s^-1^; ∼ 0.98 mV s^-1^ per 1°C change in temperature of SD) after cold acclimation (37). Interestingly, the TPP recovery slope after SD did not differ between groups; however, this might be due to recoveries occurring at very similar temperatures.

Using anoxia to disentangle the drivers of variation in SD, we observed a ∼ 0.20 mV change in TPP amplitude per 1°C change in temperature (**Fig. 5A**). This represents only ∼ 13 % of the effect of acclimation on TPP amplitude at cold-induced SD (**Fig. 3C**), meaning this difference largely reflects an effect of acclimation status. We therefore suggest that thermal acclimation must alter one or more of the factors upstream of the TPP amplitude (e.g. V_a_). Indeed, recent research posits a redistribution of electrogenic Na^+^/K^+^-ATPases to the adglial membrane following cold acclimation or heat hardening in insects (37, 68). Similarly, inhibition of the Na^+^/K^+^-ATPase leads to a lowered amplitude in the direct current field potential (similar to the TPP) in mice without changing the degree of K^+^ balance disruption (69). Furthermore, cold acclimation and adaptation confer reduced K^+^ balance disruption during SD (17), while rapid cold hardening has the opposite effect (70), meaning that electrochemical gradients likely also play a role. Interestingly, the ∼ 0.20 mV per 1°C effect we observe here closely resembles the effect of temperature on equilibrium potentials calculated by the Nernst and Goldman-Hodgkin-Katz equations (∼ 0.28 mV and ∼ 0.2 mV per 1°C change at 25°C, respectively) (see Overgaard and MacMillan (20)). Thus, temperature likely affects SD amplitude through direct thermodynamic effects on the electrochemical potentials of the transperineurial glia. Regardless, there is a significant discrepancy between the TPP amplitudes of cold- and anoxia-induced SD which might suggest that different mechanisms trigger the event, for example through different degrees of ion redistribution during SD (70) or differential effects on the basolateral membrane (V_b_) which also influences the TPP.

The direct effect of temperature on the onset slope of TPP was estimated at ∼ 0.57 mV s^-1^ per 1°C change in temperature. This represents ∼ 58% of the difference observed between the acclimation groups during cold-induced SD (**Fig. 5B** and **Fig. 3D**, respectively). This is in stark contrast to the ∼ 13% of the TPP amplitude effect, and is counter to the supposed auto-correlative nature of these two parameters (i.e. with similar timing a larger amplitude should lead to a steeper slope). However, a similar independence between these two factors has been seen in mammalian SD (71). The route through which ions and water move during SD remains a topic of debate and inferences therefore remain speculative (3, 30), however, the rate at which the TPP changes during the onset of SD must, at least partially, represent the rate at which the electrochemical gradients change, and by extension how fast ions move. The pre-SD and SD states can be described by a series of thermodynamic and electrochemical models (30), however, the thermal sensitivity of the SD onset slope is higher (Q_10_ ∼ 1.6) than what would generally be expected from ion channel mechanics (72). Thus, it seems likely that the large, progressive disruption of ion gradients occurring during SD involves an active biochemical process, which slows at low temperature, or that it is modulated by an active process which is temperature sensitive. That being said, diffusion of Ca^2+^ through the cytosol has been shown to have a relatively high Q_10_ of 2.04 (73), and so does conductance through some Ca^2+^ channels (72). Not surprisingly, a key role of Na^+^/K^+^-ATPase dysfunction has been suggested, specifically its transformation into a large non-specific conductance channel (74), and it is therefore tempting to infer that the adaptive changes to Na^+^/K^+^- ATPase activity observed for cold-acclimated flies may play a part (37). Regardless, it is clear that the progression of transmembrane ion balance disruption is slowed at low temperature; however, until a mechanism of disruption, or route of ion or water movement has been identified, further discussion regarding the mechanisms underlying our observations will remain speculative.

The TPP recovery slope did not differ between acclimation groups, likely because the temperature at which the slope was steepest was similar (**Fig. 3E,F**). In the anoxia experiments we found a non-linear relationship between recovery slope and temperature that resembled a thermal performance curve (**Fig. 5C**), which makes obtaining a comparable measure of thermal sensitivity challenging. However, calculations based on mean thermal sensitivities and maximum recovery rates performed at relevant temperatures (16.6 and 17.6°C for cold- and warm-acclimated flies, respectively; see **Fig. 3F**) resulted in a ∼ 0.16 mV s^-1^ change per 1°C difference in temperature, which represents ∼ 40 % of the effect seen at cold-induced SD. Re-establishing electrochemical gradients after SD is in large part driven by the activity of the Na^+^/K^+^-ATPase (43, 49), and an effect of temperature is therefore not surprising. In support of this, the study by Armstrong, Rodríguez and Robertson (70) found a strong effect of temperature on K^+^ concentration recovery rates after an anoxia-induced SD in fruit flies, and a faster rate of recovery in more cold-tolerant flies after cold-induced SD. Lastly, it is highly unlikely that changes to Na^+^/K^+^-ATPase activity alone explain differences in recovery from SD. Indeed, TPP recovery is affected by a wide range of other ionoregulatory processes, for example ligand-gated K^+^ channels and V-type H^+^-ATPase activity, all of which are temperature-sensitive to some extent (10, 43, 49).

### Functional aspects of temperature effects on the pre- and post-anoxic events

The smaller shifts in the TPP waveform that occur immediately before and after the SD have been categorized and linked to distinct physiological function. We did not observe either the pre- or the post- SD events during cold-induced SD. By contrast, they were always present when the SD was triggered by anoxia (**Fig. 6**). Interestingly, pre-SD events have been observed in locust preparations during rapid cooling events (9), indicating species-specific neural responses to stressful cold.

Regardless, the pre-anoxic positivity has been hypothesized to stem from a change in activity of electrogenic ATPases (42). In support of this, we found a positive effect of temperature on the amplitude and a negative effect on the width of the event. The latter was amplified by cold acclimation, which might indicate an energy-dependent process that works slower (lowered amplitude) and takes longer to deplete energy sources (duration) at low temperature (**Fig. 6A-D**). Interestingly, this results in an event whose overall magnitude is unaffected by temperature, yet increased by cold acclimation (**Fig. 6C,D**), indicating that cold-acclimated flies could have ATPases capable of continuing work as temperature is lowered. This closely matches our current understanding of the relationship between cold tolerance and Na^+^/K^+^-ATPase activity (37), and the notion that Na^+^/K^+^-ATPase activity in the adglial membrane will tend to drive the TPP upwards (creating a more positive baseline TPP, and larger pre-anoxic positivity, **Fig. 3B**). The improved ability to prevent SD may also be linked to Na^+^/K^+^-ATPase trafficking to adglial or neuronal membranes (68, 75). Interestingly, a recent study showed that cold-acclimated insects were also able to maintain endocrine stimulation at low temperature (76). Thus, electrogenic ATPases in cold-acclimated flies might be able to maintain activity through continued stimulation by an endocrine factor, rather than endogenous changes to the ATPases. An alternative explanation may be found in the V-type H^+^-ATPase, for which increased activity will act to drive the TPP in the opposite direction (49). Thus, the pre-anoxic positivity could in theory relate to increased Na^+^/K^+^-ATPase activity, reduced V-type H^+^-ATPase activity or a combination of both. Nonetheless, the strongest evidence currently available suggests that the thermally insensitive pre-anoxic event after cold acclimation and its increased magnitude relates to an improved neural or glial ionoregulatory capacity (e.g. by the Na^+^/K^+^-ATPase).

The post-anoxic negativity has been linked to reactivation of V-type H^+^-ATPases, which work to re-establish disrupted proton gradients across neural and adglial membranes after reoxygenation (49), and involvement of an electrogenic ATPase is directly supported by the temperature-sensitive nature of the event in our data. Furthermore, our findings might indicate that neural V-type H^+^- ATPase activity is unaffected by thermal acclimation (i.e. no effects of acclimation on its characteristics). However, the magnitude of the post-anoxic negativity is masked by Na^+^/K^+^- ATPase activity, such that increased Na^+^/K^+^-ATPase activity will tend to shorten the negativity and lower its amplitude (see (49)). Changes to neural Na^+^/K^+^-ATPase activity in relation to temperature and thermal acclimation are well-established (37, 75) and we therefore caution against any inference based our these findings here alone. That being said, it seems likely that changes to the V- type H^+^-ATPase are involved in modulating the effects of temperature and thermal acclimation on the SD threshold.

### Cold acclimation lowers the thermal sensitivity of spreading depolarization propagation

A key aspect of the SD is the ability to propagate from the point of initially compromised function and into naïve neural tissue (5), and this is also true for *Drosophila* (42). We found that the propagation velocity was highly temperature-sensitive and resembled an exponential function, and that the propagation velocity in cold-acclimated flies was less thermally sensitive that in warm- acclimated flies (**Fig. 7**). Furthermore, our estimates at room temperature were similar to previous estimates for *Drosophila melanogaster* (13, 42).

Effects of temperature on propagation velocity of SD have been observed previously in tissues of both endotherms (chicken and monkey (46, 77)), and ectotherms (frog and skate (78, 79)), and in most cases the relationship largely appears exponential with a Q_10_ around 2-3, which matches those obtained for fruit flies here (2.1 and 3.0 for cold- and warm-acclimated flies, respectively).

The physiological mechanism underlying the ability for SD to propagate across tissue, like its trigger mechanism, remains a topic of debate, however, the thermal dependency of the propagation velocity and the degree of thermal sensitivity (i.e. Q_10_) strongly indicates the involvement of an active biochemical component to the spread (80, 81). The current consensus model on the mechanism propagation is that of a reaction-diffusion process, where the reaction (i.e. SD) generates an activator, which subsequently diffuses and triggers the reaction in adjacent tissue (3, 82). Our findings support this model, as we find strong evidence for an active biochemical component, which we assume relates to the reaction part of the model.

The propagation velocity of a SD is affected by a multitude of factors (see (5)), however, this is the first time the impact of thermal history on this factor has been investigated. Given that the mechanism of the spread remains unknown (but must involve an active component, see above), any inferences on the adaptive or maladaptive nature of reduced spreading velocity in cold-acclimated flies would be speculative. Nonetheless, it is interesting to note that despite the different thermal sensitivities, the rate of propagation at the temperatures that cause SD *per se* are likely similar at ∼ 0.6-0.7 mm min^-1^ (based on extrapolation). Similarly, the overall relationship with temperature closely resembles that of the activity of the Na^+^/K^+^-ATPase: Warm-acclimated flies have higher propagation velocities and Na^+^/K^+^-ATPase activities at higher temperatures which are more heavily reduced at lower temperature when compared to cold-acclimated flies, and with a cross-over at ∼ 15°C (**Fig. 7**, and Fig. 5 in (37)). As mentioned above, the thermal sensitivity of the propagation velocity strongly suggests the involvement of an active biochemical process, many of which have improved function at low temperature after cold acclimation or adaptation or display reduced thermal sensitivity (10, 83, 84). This is supported by the reduced Q_10_ of the cold-acclimated flies (2.12) compared to their warm-acclimated conspecifics (2.98). Similar support is found after an Arrhenius transformation of the datasets which shows a reduction in activation energy from 80.8 kJ mol^-1^ (0.84 eV) to 55.9 kJ mol^-1^ (0.58 eV) after cold-acclimation. These values are similar to those found for biochemical processes in other ectotherms (85) and are also similar to those found for the Na^+^/K^+^-ATPase previously in the brains of fruit fly acclimated to the same temperatures (80.4 kJ mol^-1^ and 59.0 kJ mol^-1^) (37) and mammalian tissues (although these vary considerable based on experimental conditions (86, 87)). Given the putative role of the Na^+^/K^+^-ATPase and its failure in triggering the SD itself it seems less speculative to implicate it in SD propagation, however, further research is needed to elucidate the mechanisms of propagation and to dissect the causal links between propagation velocity, temperature, and thermal acclimation.

## Conclusion

In summary, we demonstrate that thermal acclimation changes the temperature-threshold for cold-induced SD as well as the associated waveform of the transperineurial potential, and that at least part of this effect on the waveform is driven by temperature *per se* (see **Fig. 8**). Subsequently, we use specific parameters of the waveform to speculate on potential mechanisms underlying variation in the threshold for cold- and anoxia-induced SD, while also generating testable hypotheses.

**Figure 8.**
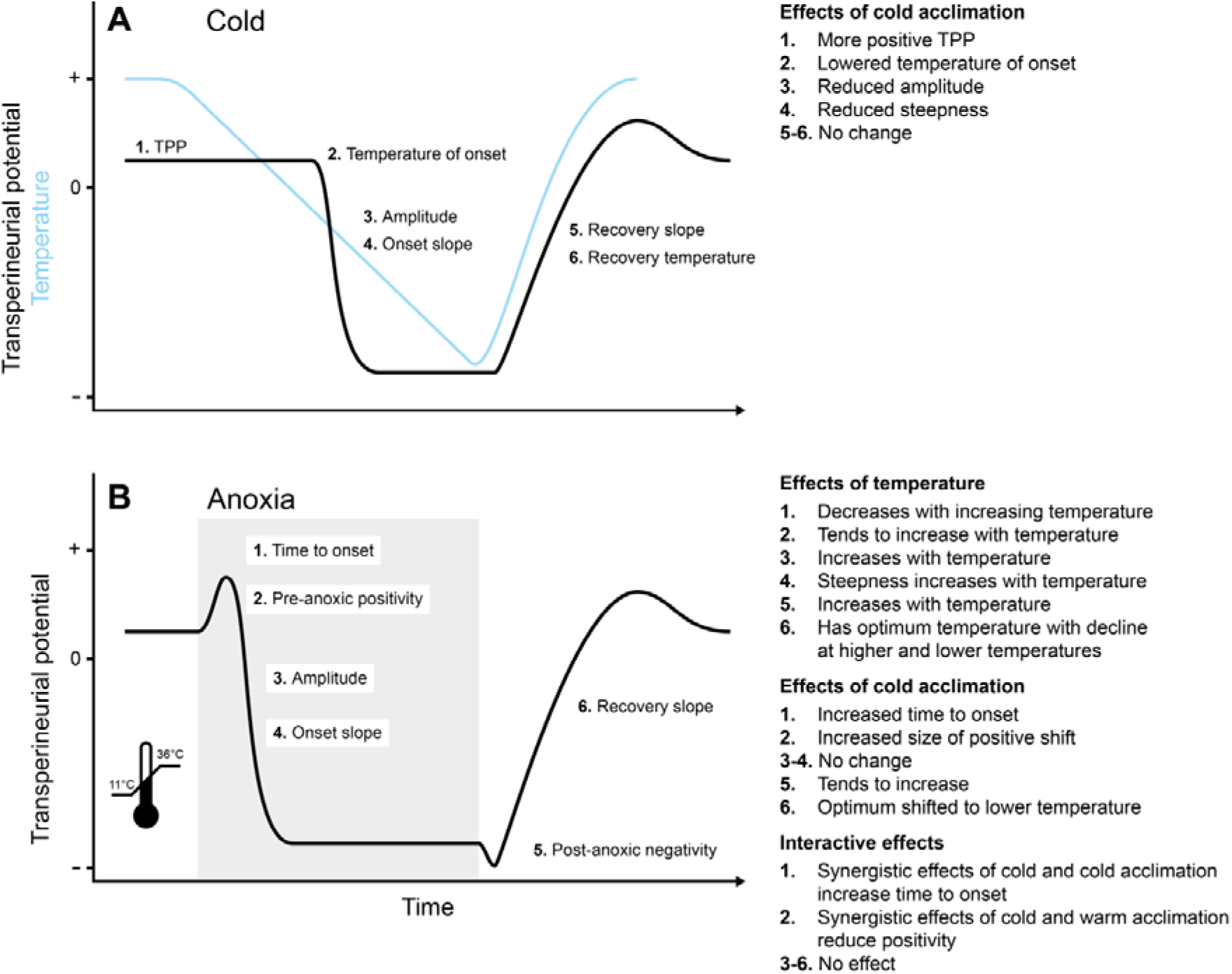
Summary of the isolated and combined effects of temperature and thermal acclimation on the threshold for cold- and anoxia-induced spreading depolarization and their associated transperineurial waveforms. (A) Stressful cold lead to spreading depolarization and cold acclimation lowers the temperature of onset while also eliciting specific changes to the associated transperineurial waveform (A, right, **see Fig. 3**). To investigate if these changes to the waveform were due to acclimation or if they were a consequence of the spreading depolarization occurring at different temperatures we quantified the waveform at different temperatures with anoxia (B). Anoxia (shaded area in B) led to a slightly different transperineurial waveform, of which the pre- and post-anoxic shifts were the most notable. Nonetheless, the effects of temperature and thermal acclimation, and their interaction, were estimated (B, right, **see Figs. 4-6**). This revealed that a direct effect of temperature was present for all parameters, which meant that this effect likely played a role (of varying magnitude, see main text) in the differences observed between acclimation groups at low temperature. Specifically, temperature played a small, likely thermodynamic, role in the amplitude of the transperineurial waveform (A3 and B3) while it explained a large part of the differences seen in the slope of onset (A4 and B4) and recovery (A5 and B6).

Specifically, we highlight potential roles for adaptive changes to the Na^+^/K^+^-ATPase, V-type H^+^- ATPase, and K^+^ conductance. We also note distinct differences in the waveform when SD was induced with cold and anoxia. The rate at which a SD propagated across the brain was temperature-sensitive and this sensitivity was reduced by cold acclimation, suggesting the involvement of an active biochemical process. Further research is needed to elucidate the physiological mechanisms underlying improved abiotic stress tolerance of the CNS, as limited by SD, and the role of changes to thermal sensitivity of the propagation velocity.

## Acknowledgements

We would like to extend our gratitude to Jeff Dawson (Department of Biology, Carleton University) for lending us the Neuroprobe amplifier which allowed us to measure the spreading depolarization propagation velocity.

## Author contributions

The study was conceived and designed by MKA. MKA performed the experiments, analyzed the data, and wrote first draft. All authors made edits to the manuscript and approved the final version of the manuscript.

## Data availability

All data has been made available to reviewers during the review process and will be uploaded as supplementary material upon acceptance and publication.

---

**Fig. S1.**
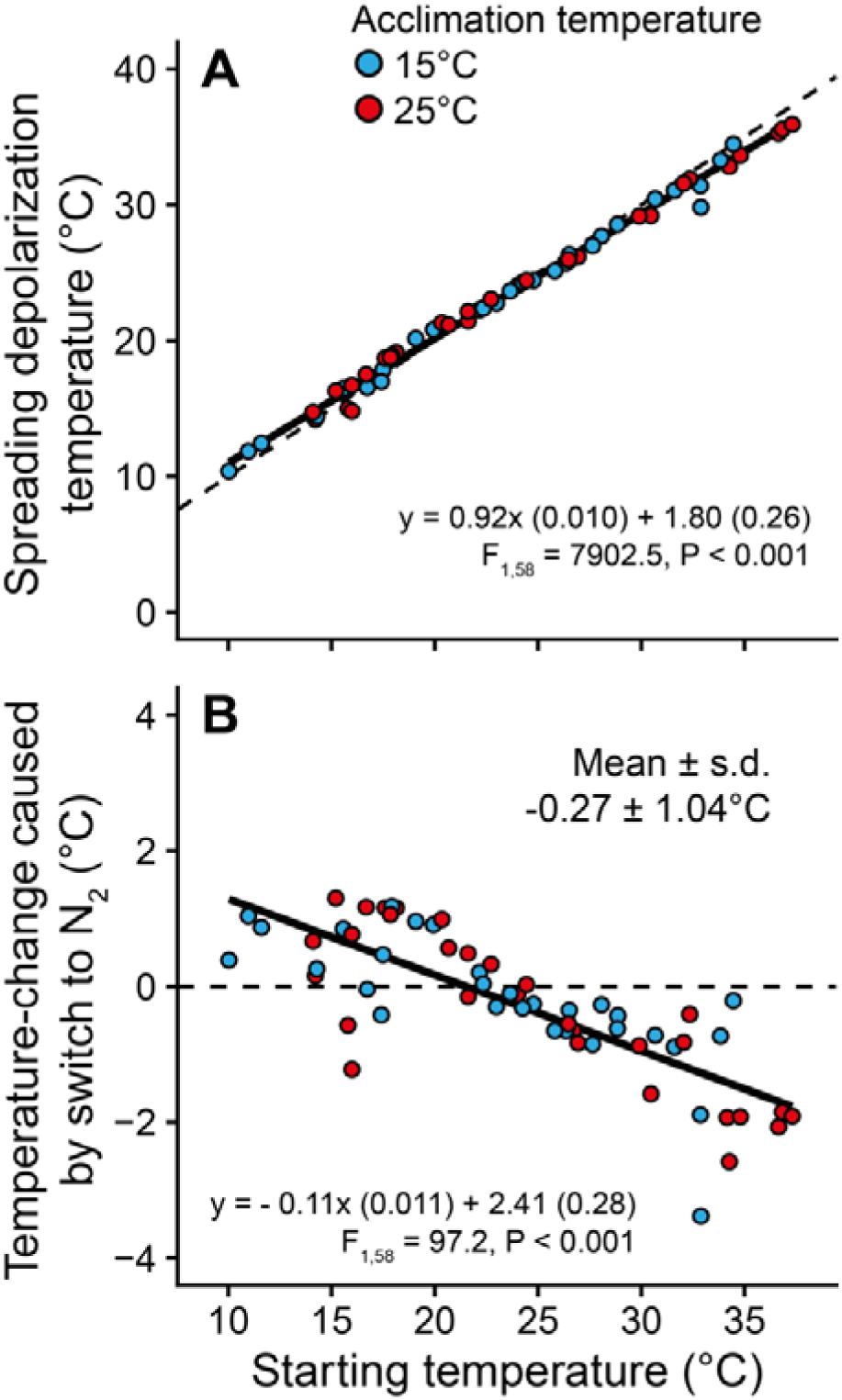
Effect of switching from air to N_2_ exposure on the temperature measured by the thermocouple next to the fly head in experiments on the effects of temperature on the TPP waveform during anoxia-induced spreading depolarization. During experiments designed to investigate the effect of thermal acclimation and temperature on the changes to the TPP during anoxia-induced spreading depolarization the switch from atmospheric air to N_2_ gas resulted in small temperature changes. This meant that the starting temperature often differed slightly from the temperature recorded at the onset of the spreading depolarization. (A) The relationship between these two temperatures (i.e. starting temperature and temperature recorded at the onset of the spreading depolarization) was linear and deviated only little from the line of unity (dashed line). (B) The effect of the switch in gas was generally small (-0.27 ± 1.04°C, mean ± standard deviation) and mostly changed the temperature towards that of the room (∼ 23°C). A statistical analysis (one-sample t test comparing all values to 0) revealed a borderline significant effect of the change (t_59_ = - 2.0, P = 0.051), nonetheless, there is clear effect of starting temperature on the magnitude of the temperature change. Numbers in parenthesis represent the error on the parameter estimates by the linear models.

## References

1. Robertson RM, Dawson-Scully KD, and Andrew RD. Neural shutdown under stress: an evolutionary perspective on spreading depolarization. Journal of Neurophysiology 123: 885–895, 2020.

2. Leao AA. Spreading depression of activity in the cerebral cortex. Journal of neurophysiology 7: 359–390, 1944.

3. Andrew RD, Hartings JA, Ayata C, Brennan K, Dawson-Scully KD, Farkas E, Herreras O, Kirov S, Müller M, and Ollen-Bittle N. The critical role of spreading depolarizations in early brain injury: consensus and contention. Neurocritical Care 1–19, 2022.

4. Rodgers CI, Armstrong GA, and Robertson RM. Coma in response to environmental stress in the locust: a model for cortical spreading depression. Journal of Insect Physiology 56: 980–990, 2010.

5. Pietrobon D, and Moskowitz MA. Chaos and commotion in the wake of cortical spreading depression and spreading depolarizations. Nature Reviews Neuroscience 15: 379, 2014.

6. Dreier JP, and Reiffurth C. The stroke-migraine depolarization continuum. Neuron 86: 902–922, 2015.

7. Shuttleworth CW, Andrew RD, Akbari Y, Ayata C, Balu R, Brennan K, Boutelle M, Carlson AP, Dreier JP, Fabricius M, Farkas E, Foreman B, Helbok R, Henninger N, Jewell SL, Jones SC, Kirov SA, Lindquist BE, Maciel CB, Okonkwo D, Reinhart KM, Robertson RM, Rosenthal ES, Watanabe T, and Hartings JA. Which Spreading Depolarizations Are Deleterious To Brain Tissue? In: Neurocritical care Springer, 2019, p. 1-6.

8. Jørgensen LB, Robertson RM, and Overgaard J. Neural dysfunction correlates with heat coma and CT_max_ in *Drosophila* but does not set the boundaries for heat stress survival. Journal of Experimental Biology 2020.

9. Robertson RM, Spong KE, and Srithiphaphirom P. Chill coma in the locust, *Locusta migratoria*, is initiated by spreading depolarization in the central nervous system. Scientific reports 7: 10297, 2017.

10. Robertson RM, MacMillan HA, and Andersen MK. A cold and quiet brain: mechanisms of insect CNS arrest at low temperatures. Current Opinion in Insect Science 101055, 2023.

11. Andersen MK, Willot Q, and MacMillan HA. A neurophysiological limit and its biogeographic correlations: cold-induced spreading depolarization in tropical butterflies. Journal of Experimental Biology 226: 2023.

12. Andreassen AH, Hall P, Khatibzadeh P, Jutfelt F, and Kermen F. Brain dysfunction during warming is linked to oxygen limitation in larval zebrafish. Proceedings of the National Academy of Sciences 119: e2207052119, 2022.

13. Spong KE, Andrew RD, and Robertson RM. Mechanisms of spreading depolarization in vertebrate and insect central nervous systems. Journal of Neurophysiology 116: 1117–1127, 2016.

14. MacMillan HA, and Sinclair BJ. Mechanisms underlying insect chill-coma. Journal of Insect Physiology 57: 12–20, 2011.

15. Semper C. The natural conditions of existence as they affect animal life. Kegan Paul, Trench & Company, 1883.

16. Mellanby K. Low temperature and insect activity. Proceedings of the Royal Society of London Series B, Biological Sciences 473–487, 1939.

17. Andersen MK, Jensen NJS, Robertson RM, and Overgaard J. Central nervous system shutdown underlies acute cold tolerance in tropical and temperate *Drosophila* species. Journal of Experimental Biology 221: 2018.

18. Andersen MK, and Overgaard J. The central nervous system and muscular system play different roles for chill coma onset and recovery in insects. Comparative Biochemistry and Physiology Part A: Molecular & Integrative Physiology 233: 10–16, 2019.

19. Hazell SP, and Bale JS. Low temperature thresholds: Are chill coma and CT_min_ synonymous? Journal of Insect Physiology 57: 1085–1089, 2011.

20. Overgaard J, and MacMillan HA. The integrative physiology of insect chill tolerance. Annual Review of Physiology 79: 187–208, 2017.

21. Addo-Bediako A, Chown SL, and Gaston KJ. Thermal tolerance, climatic variability and latitude. Proceedings of the Royal Society of London B: Biological Sciences 267: 739–745, 2000.

22. Weaving H, Terblanche JS, Pottier P, and English S. Meta-analysis reveals weak but pervasive plasticity in insect thermal limits. Nature communications 13: 5292, 2022.

23. Kellermann V, Loeschcke V, Hoffmann A, Kristensen T, Flojgaard C, David J, Svenning J-C, and Overgaard J. Phylogenetic constraints in key functional traits behind species’ climate niches: patterns of desiccation and cold resistance across 95 *Drosophila* species. Evolution; international journal of organic evolution [P] 66: 3377–3389, 2012.

24. Kimura MT. Cold and heat tolerance of drosophilid flies with reference to their latitudinal distributions. Oecologia 140: 442–449, 2004.

25. Schou MF, Mouridsen MB, Sørensen JG, and Loeschcke V. Linear reaction norms of thermal limits in *Drosophila*: predictable plasticity in cold but not in heat tolerance. Functional Ecology 31: 934–945, 2017.

26. Overgaard J, Kristensen TN, Mitchell KA, and Hoffmann AA. Thermal tolerance in widespread and tropical *Drosophila* species: does phenotypic plasticity increase with latitude? The American Naturalist 178: S80–S96, 2011.

27. Ransberry VE, MacMillan HA, and Sinclair BJ. The relationship between chill-coma onset and recovery at the extremes of the thermal window of *Drosophila melanogaster*. Physiological and Biochemical Zoology 84: 553–559, 2011.

28. MacMillan HA, Andersen JL, Loeschcke V, and Overgaard J. Sodium distribution predicts the chill tolerance of *Drosophila melanogaster* raised in different thermal conditions. *American Journal of Physiology-Regulatory*, Integrative and Comparative Physiology 308: R823–R831, 2015.

29. Cheslock A, Andersen MK, and MacMillan HA. Thermal acclimation alters Na^+^/K^+^-ATPase activity in a tissue-specific manner in *Drosophila melanogaster*. Comparative Biochemistry and Physiology Part A: Molecular & Integrative Physiology 110934, 2021.

30. Lemale CL, Lückl J, Horst V, Reiffurth C, Major S, Hecht N, Woitzik J, and Dreier JP. Migraine Aura, Transient Ischemic Attacks, Stroke, and Dying of the Brain Share the Same Key Pathophysiological Process in Neurons Driven by Gibbs–Donnan Forces, Namely Spreading Depolarization. Frontiers in Cellular Neuroscience 16: 40, 2022.

31. Schofield P, and Treherne J. Localization of the blood-brain barrier of an insect: electrical model and analysis. Journal of experimental biology 109: 319–331, 1984.

32. Spong KE, Chin B, Witiuk KL, and Robertson RM. Cell swelling increases the severity of spreading depression in *Locusta migratoria*. Journal of neurophysiology 114: 3111–3120, 2015.

33. Srithiphaphirom P, and Robertson RM. Rapid cold hardening delays the onset of anoxia-induced coma via an octopaminergic pathway in Locusta migratoria. Journal of Insect Physiology 137: 104360, 2022.

34. Andersen MK, Willot Q, and MacMillan HA. Tropical butterflies lose central nervous system function in the cold from a spreading depolarization event. bioRxiv 2023.2005. 2031.543057, 2023.

35. Terai H, Gwedela MNV, Kawakami K, and Aizawa H. Electrophysiological and pharmacological characterization of spreading depolarization in the adult zebrafish tectum. Journal of Neurophysiology 126: 1934–1942, 2021.

36. Schofield P. Oscillations of glial membrane potential in a localized region of the blood-brain barrier of an insect. Journal of experimental biology 148: 335–351, 1990.

37. Andersen MK, Robertson RM, and MacMillan HA. Plasticity in Na^+^/K^+^-ATPase thermal kinetics drives variation in the temperature of cold-induced neural shutdown of adult *Drosophila melanogaster*. Journal of Experimental Biology 2022.

38. Goldman DE. Potential, impedance, and rectification in membranes. The Journal of General Physiology 27: 37–60, 1943.

39. Hodgkin AL, and Katz B. The effect of sodium ions on the electrical activity of the giant axon of the squid. The Journal of physiology 108: 37, 1949.

40. Fraser JA, and Huang CLH. A quantitative analysis of cell volume and resting potential determination and regulation in excitable cells. The Journal of Physiology 559: 459–478, 2004.

41. Bayley JS, Overgaard J, and Pedersen TH. Quantitative model analysis of the resting membrane potential in insect skeletal muscle: Implications for low temperature tolerance. Comparative Biochemistry and Physiology Part A: Molecular & Integrative Physiology 257: 110970, 2021.

42. Spong KE, Rodríguez EC, and Robertson RM. Spreading depolarization in the brain of *Drosophila* is induced by inhibition of the Na^+^/K^+^-ATPase and mitigated by a decrease in activity of protein kinase G. Am Physiological Soc, 2016.

43. Van Dusen RA, Shuster-Hyman H, and Robertson RM. Inhibition of ATP-sensitive potassium channels exacerbates anoxic coma in *Locusta migratoria*. Journal of Neurophysiology 124: 1754–1765, 2020.

44. Rodgers CI, Armstrong GA, Shoemaker KL, LaBrie JD, Moyes CD, and Robertson RM. Stress preconditioning of spreading depression in the locust CNS. PloS one 2: e1366, 2007.

45. Somjen GG. Mechanisms of spreading depression and hypoxic spreading depression-like depolarization. Physiological reviews 81: 1065–1096, 2001.

46. Martins-Ferreira H, De Oliveira Castro G, Struchiner C, and Rodrigues P. Circling spreading depression in isolated chick retina. Journal of Neurophysiology 37: 773–784, 1974.

47. Marshall KE, and Sinclair BJ. Repeated stress exposure results in a survival– reproduction trade-off in *Drosophila melanogaster*. Proceedings of the Royal Society B: Biological Sciences 277: 963–969, 2010.

48. Nilson TL, Sinclair BJ, and Roberts SP. The effects of carbon dioxide anesthesia and anoxia on rapid cold-hardening and chill coma recovery in *Drosophila melanogaster*. Journal of insect physiology 52: 1027–1033, 2006.

49. Robertson RM, and Van Dusen RA. Motor patterning, ion regulation and spreading depolarization during CNS shutdown induced by experimental anoxia in *Locusta migratoria*. Comparative Biochemistry and Physiology Part A: Molecular & Integrative Physiology 260: 111022, 2021.

50. Rodríguez EC, and Robertson RM. Protective effect of hypothermia on brain potassium homeostasis during repetitive anoxia in *Drosophila melanogaster*. Journal of Experimental Biology 215: 4157–4165, 2012.

51. R Core Team. R: A language and environment for statistical computing. R Foundation for Statistical Computing: See http://www.R-project.org/, 2023.

52. Birk M. respirometry: Tools for conducting and analyzing respirometry experiments. *R package, version 07 0, URL:* http://cran *r-project org/package= respirometry* 2018.

53. Dreier JP. The role of spreading depression, spreading depolarization and spreading ischemia in neurological disease. Nature medicine 17: 439–447, 2011.

54. Benasayag-Meszaros R, Risley MG, Hernandez P, Fendrich M, and Dawson-Scully K. Pushing the limit: examining factors that affect anoxia tolerance in a single genotype of adult D. melanogaster. Scientific reports 5: 1–5, 2015.

55. Storey KB, and Storey JM. Oxygen: Stress and adaptation in cold-hardy insects. Low temperature biology of insects 141–165, 2010.

56. Lutz PL, Nilsson GE, and Prentice HM. The brain without oxygen: causes of failure-physiological and molecular mechanisms for survival. Springer Science & Business Media, 2003.

57. Nilsson GE, and Lutz PL. Anoxia tolerant brains. Journal of Cerebral Blood Flow & Metabolism 24: 475–486, 2004.

58. MacMillan HA, Knee JM, Dennis AB, Udaka H, Marshall KE, Merritt TJ, and Sinclair BJ. Cold acclimation wholly reorganizes the *Drosophila melanogaster* transcriptome and metabolome. Scientific Reports 6: 28999, 2016.

59. Colinet H, Larvor V, Laparie M, and Renault D. Exploring the plastic response to cold acclimation through metabolomics. Functional Ecology 26: 711–722, 2012.

60. Berrigan D, and Partridge L. Influence of temperature and activity on the metabolic rate of adult *Drosophila melanogaster*. Comparative Biochemistry and Physiology Part A: Physiology 118: 1301–1307, 1997.

61. Alton LA, Condon C, White CR, and Angilletta Jr MJ. Colder environments did not select for a faster metabolism during experimental evolution of *Drosophila melanogaster*. Evolution 71: 145–152, 2017.

62. Messamah B, Kellermann V, Malte H, Loeschcke V, and Overgaard J. Metabolic cold adaptation contributes little to the interspecific variation in metabolic rates of 65 species of Drosophilidae. Journal of insect physiology 98: 309–316, 2017.

63. Hochachka P. Defense strategies against hypoxia and hypothermia. Science 231: 234–241, 1986.

64. Hochachka P, Buck L, Doll C, and Land S. Unifying theory of hypoxia tolerance: molecular/metabolic defense and rescue mechanisms for surviving oxygen lack. Proceedings of the National Academy of Sciences 93: 9493–9498, 1996.

65. Boutilier RG. Mechanisms of cell survival in hypoxia and hypothermia. Journal of Experimental Biology 204: 3171–3181, 2001.

66. Robertson RM. The origin of the ‘channel arrest’ hypothesis. Journal of Experimental Biology 220: 1747–1748, 2017.

67. Schofield P, and Treherne J. Octopamine reduces potassium permeability of the glia that form the insect blood-brain barrier. Brain research 360: 344–348, 1985.

68. Hou N, Armstrong GA, Chakraborty-Chatterjee M, Sokolowski MB, and Robertson RM. Na^+^-K^+^-ATPase trafficking induced by heat shock pretreatment correlates with increased resistance to anoxia in locusts. Journal of neurophysiology 112: 814–823, 2014.

69. Reiffurth C, Alam M, Zahedi-Khorasani M, Major S, and Dreier JP. Na^+^/K^+^- ATPase α isoform deficiency results in distinct spreading depolarization phenotypes. Journal of Cerebral Blood Flow & Metabolism 40: 622–638, 2020.

70. Armstrong GA, Rodríguez EC, and Robertson RM. Cold hardening modulates K^+^ homeostasis in the brain of *Drosophila* melanogaster during chill coma. Journal of Insect Physiology 58: 1511–1516, 2012.

71. Menyhárt Á, Makra P, Szepes BÉ, Tóth OM, Hertelendy P, Bari F, and Farkas E. High incidence of adverse cerebral blood flow responses to spreading depolarization in the aged ischemic rat brain. Neurobiology of aging 36: 3269–3277, 2015.

72. Janssen R. Thermal influences on nervous system function. Neuroscience & Biobehavioral Reviews 16: 399–413, 1992.

73. Sidell BD, and Hazel JR. Temperature affects the diffusion of small molecules through cytosol of fish muscle. Journal of Experimental Biology 129: 191–203, 1987.

74. Brisson CD, Hsieh Y-T, Kim D, Jin AY, and Andrew RD. Brainstem neurons survive the identical ischemic stress that kills higher neurons: insight to the persistent vegetative state. PloS one 9: e96585, 2014.

75. Robertson RM, and Moyes CD. Rapid cold hardening increases axonal Na^+^/K^+^- ATPase activity and enhances performance of a visual motion detection circuit in *Locusta migratoria*. Journal of Experimental Biology 225: jeb244097, 2022.

76. Gerber L, Kresse J-C, Šimek P, Berková P, and Overgaard J. Cold acclimation preserves hindgut reabsorption capacity at low temperature in a chill-susceptible insect, *Locusta migratoria*. Comparative Biochemistry and Physiology Part A: Molecular & Integrative Physiology 252: 110850, 2021.

77. Marshall W, Essig C, and Dubroff S. Relation of temperature of cerebral cortex to spreading depression of Leão. Journal of Neurophysiology 14: 153–166, 1951.

78. Martins-Ferreira H, and Carmo Rd. Retinal spreading depression and the extracellular milieu. Canadian Journal of Physiology and Pharmacology 65: 1092–1098, 1987.

79. Young W. Spreading depression in elasmobranch cerebellum. Brain research 199: 113–126, 1980.

80. Cossins A. Temperature biology of animals. Springer Science & Business Media, 2012.

81. Somero GN. Adaptation of enzymes to temperature: searching for basic “strategies”. Comparative Biochemistry and Physiology Part B: Biochemistry and Molecular Biology 139: 321–333, 2004.

82. Zandt B-J, ten Haken B, and van Putten MJ. Diffusing substances during spreading depolarization: analytical expressions for propagation speed, triggering, and concentration time courses. Journal of Neuroscience 33: 5915-5923, 2013.

83. Hochachka PW, and Somero GN. Strategies of biochemical adaptation. 1973.

84. Hochachka PW, and Somero GN. Biochemical adaptation: mechanism and process in physiological evolution. Oxford university press, 2002.

85. Jørgensen LB, Ørsted M, Malte H, Wang T, and Overgaard J. Extreme escalation of heat failure rates in ectotherms with global warming. Nature 611: 93–98, 2022.

86. Charnock J, Almeida A, and To R. Temperature-activity relationships of cation activation and ouabain inhibition of (Na++ K+)-ATPase. Archives of Biochemistry and Biophysics 167: 480–487, 1975.

87. Swann AC. (Na+, K+)-ATPase of mammalian brain: Effects of temperature on cation and ATP interactions regulating phosphatase activity. Archives of Biochemistry and Biophysics 221: 148–157, 1983.

